# Dynamic substrate preferences and predicted metabolic properties of a simple microbial consortium

**DOI:** 10.1101/053777

**Authors:** Onur Erbilgin, Benjamin P. Bowen, Suzanne M. Kosina, Stefan Jenkins, Rebecca K. Lau, Trent R. Northen

**Affiliations:** Environmental Genomics and Systems Biology Division, Lawrence Berkeley National Laboratory, 1 Cyclotron Road, Berkeley, Berkeley, CA 94720; Joint Genome Institute, 2800 Mitchell Dr. Walnut Creek, CA, 94598 Joint Genome Institute, 2800 Mitchell Dr. Walnut Creek, CA, 94598

## Abstract

Microorganisms are typically found as complex microbial communities that altogether govern global biogeochemical cycles. Microbes have developed highly regulated metabolic capabilities to efficiently use available substrates including preferential substrate usage that can result in diauxic shifts. This and other metabolic behaviors have been discovered in studies of microbes in monoculture when grown on low-complexity (*e.g.* two-component) mixtures of substrates, however, little is known about how species partition environmental substrates through substrate competition in more complex substrate mixtures. Here we use exometabolomic profiling to examine the time-varying substrate depletion from a mixture of 19 amino acids and glucose by two *Pseudomonads* and one *Bacillus* species isolated from ground water. We examine if the first substrates depleted result in maximal growth rate, or relate to growth medium or biomass composition and find surprisingly few correlations. Patterns of substrate depletion are modeled, and these models are used to examine if substrate usage preferences and substrate depletion kinetics of three microbial isolates can be used to predict the metabolism of the pooled isolates in co-culture. We find that most of the substrates fit the model predictions, indicating that the microbes are not altering their behaviors for these substrates in the presence of competitors. Glucose and histidine were depleted more slowly than predicted, while proline, glycine, glutamate, lysine, and arginine were all consumed significantly faster; these compounds highlight substrates that could be involved in species-species interactions within the consortium.

**Author Contributions:** OE, TRN conceived and designed the experiments

OE, BPB, SMK, SJ, RL performed the experiments

OE, BPB, SJ, TRN analyzed the data

OE, TRN wrote the manuscript

TRN contributed materials and analysis tools

## Introduction

Microbial communities drive elemental cycling Ref.^1^, such as the carbon cycle ^2^ and the nitrogen cycle ^3^. We are also learning that they are critical for the health of their host plant ^4^ or animal ^5^. In both cases the net microbial metabolic processes are of particular interest for predicting nutrient cycles ^6^,^7^ and harnessing microbes to improve host health ^8^.The environments in which microbial communities live often contain complex mixtures of substrates, and understanding how microbial communities partition these substrates is central to predicting community metabolism and developing interventions that alter community structure and/or metabolic activities.

Exometabolomics, also known as metabolic footprinting, is a powerful platform for studying how microbes and their consortia modify substrate pools, as analysis is only of the extracellular metabolites ^9^. With the development of exometabolomics pipelines, the metabolic connections between microbes have begun to be studied at a large scale and have allowed for a more comprehensive approach to monitoring the dynamic transformations of relatively complex mixtures of substrates ^9^. Some key examples include optimizing multiple steps of lignocellulose degradation ^10^,^11^, understanding metabolic interactions between species in mixed communities ^12^, and determining the ecological role of individuals within a mixed community ^13^^−^^15^. We have recently found exometabolite niche partitioning in two soil environments where sympatric microbes were found to target largely non-overlapping portions of the available substrates, thus minimizing substrate competition ^14^. These experiments were focused on the endpoint depletion of substrates by isolates, not the temporal sequence of utilization. However, the order of substrate utilization (*i.e.* substrate preferences) may further discriminate the adaptive strategies of individual organisms for common substrates.

While some work on mixed-substrate growth has been performed in continuous culture at steady state ^16^, understanding substrate usage and competition in batch cultures may have both ecological and practical applications. Many environmental processes happen with pulsed inputs: for example the release of substrates into the soil following rainfall, light-dark cycles, digestion in animals, *etc.* Additionally, some biotechnologies that use microorganisms are also batch processes, such as the large-scale fermentations of microbe-processed foods (*e.g.* cheese, wine, *etc*.). Most of these processes use mixed microbial cultures, including one-pot processes of biomass conversion to biofuels and other biosynthetic products ^17^^−^^19^. Studying the temporal substrate utilization by individuals is an important first step in developing approaches to better model these biochemical processes.

As recently shown in the pioneering work by Behrends *et al*., the kinetics of substrate depletion from a mixture of substrates can be effectively fit using a few parameters ^20^: see **Equation (1)** in **Materials and Methods**. When compared across all substrates in an environment, these parameters have great potential in providing a direct measure of an organism’s substrate preferences within that environment, effectively creating a metabolic model for the organism. Such models may be useful in classifying microorganisms for in-depth characterization of their metabolism and regulatory networks to understand the biochemical or evolutionary basis for these preferences. Furthermore, when taken into consideration with other species’ models, they may also enable the prediction of the overall net metabolism of microbial consortia by aggregating individual contributions to environmental substrate usage. Observed deviations from these predictions could help identify interspecies interactions that modulate an organism’s metabolism, *e.g.* communication and antagonism between microbes within communities.

Here we compare the temporal depletion of 20 substrates by 3 isolates and fit these data to the Behrends model (**Equation 1)**, describing their substrate preferences within this ‘environment’. We then examine if the first substrates depleted result in maximal growth rate, or relate to growth medium or biomass composition. Finally, we developed a model that simply combines the usage profiles of individual species to test if a consortium initially composed of an equal mixture of each of the three isolates consumes substrates in an identical manner to when they are grown individually, *i.e.* the presence of other microbes does not affect their substrate usage. Any deviations from this model may indicate compounds that are actively regulated. For example, if a compound is consumed significantly faster or earlier than predicted by the model, this would indicate an additional interaction between species such as synergistic or competitive growth.

## Results and Discussion

In order to determine the substrate usage profiles of individuals, we designed a defined medium composed of sufficient levels of standard vitamins, minerals, phosphate, and ammonium, and limiting levels of carbon (glucose and nineteen amino acids (see **Materials and Methods**). This medium was designed such that the species would reach stationary phase within 12 hours and every substrate could be detected in a single LC-MS run.

Bacilli and pseudomonads represent some of the most ubiquitous soil bacteria, and we selected the common soil bacterium *Bacillus cereus* for comparison with two closely related *Pseudomonas* species, *Pseudomonas lini* and *Pseudomonas baetica* (**Supplemental Figure 1**) that were isolated from groundwater; taxonomic assertions were confirmed by BLAST search results on the sequenced 16S rRNA gene. For simplicity, we will refer to the species as *Bc* (*Bacillus cereus*), *Pl*, (*Pseudomonas lini*), and *Pb* (*Pseudomonas baetica*). Each species was grown individually in the defined medium, with supernatant samples collected every hour for 12 hours, and one final time point at 26 hours.

The absolute concentrations of the 20 growth substrates were quantified at each time point, and the data were fit to a previously described model for compound depletion during microbial batch culture ^20^ (**Figure 1**, **Algorithm 1**). We observed that all compounds followed the Behrends model over the course of growth for each species, with the exception of two compounds: glycine increased over the first 5 hours of culture from all three species and then decreased logarithmically, and the methionine depletion profile for *Bc* was indeterminable due to both variance in the data and a lack of time points from 12 to 24 hours (**Supplemental File 1**). These observations corroborate previous assertions that substrate utilization by microbes in batch culture follow the shape of a logistic growth type curve ^20^^−^^22^.

**Figure 1.**
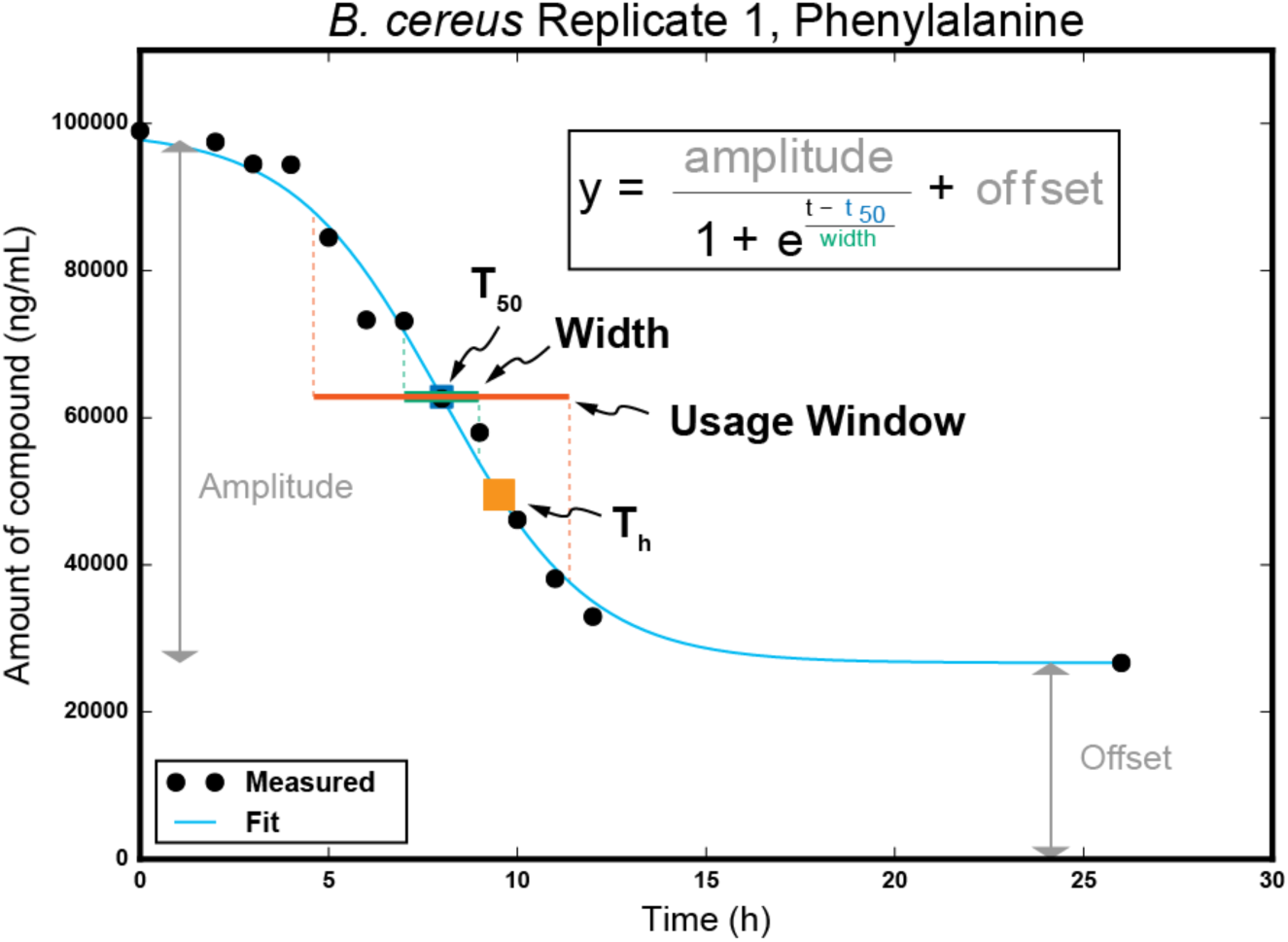
Modeling usage parameters. Example curve fitting to Behrends model (cyan). Blue square indicates the modeled T_50_ parameter of the Behrends model, or inflection point of the curve, and the width parameter of the model is depicted by the green bar centered at T_50_. The orange square represents the calculated T_h_ value, or when half of the total amount of compound has been depleted, and the red bar depicts the calculated usage window, or time when the compound is depleted from 90% to 10% of the total amount used by the species.

To examine the sequence of substrate deletion in finer detail, we used the model to calculate the time at which each species depleted half of the total amount of each compound (T_h_), and when the compound was depleted from 90% to 10% of the total amount available to the species (usage window) (**Figure 1**), and mapped them onto the growth curve of each species (**Figure 2A-C**). For *Bc*, we observed that compounds were half-depleted in three distinct groups (**Figures 2A** and **D**, dotted boxes). *Bc* initially utilized glucose, then a cluster of 13 amino acids that all had T_h_ values within 0.25 h of each other during early logarithmic growth, and finally half-depleted remaining 6 substrates in late exponential and stationary phases. Neither of the pseudomonads appeared to utilize substrates in these types of groups, but instead had a more even distribution throughout their growth curve (**Figures 2B-D**). However, the growth curve of *Pb* did show multiple growth phases (**Figure 2C**), and so compounds can be mapped to the growth phase in which they are half-depleted (**Figure 2D**). This observation is more in line with the traditional view of catabolite repression and multi-auxic growth, where a lag phase will be observed each time the organism reorganizes its metabolism to utilize different substrates ^23^.

**Figure 2. A-C).**
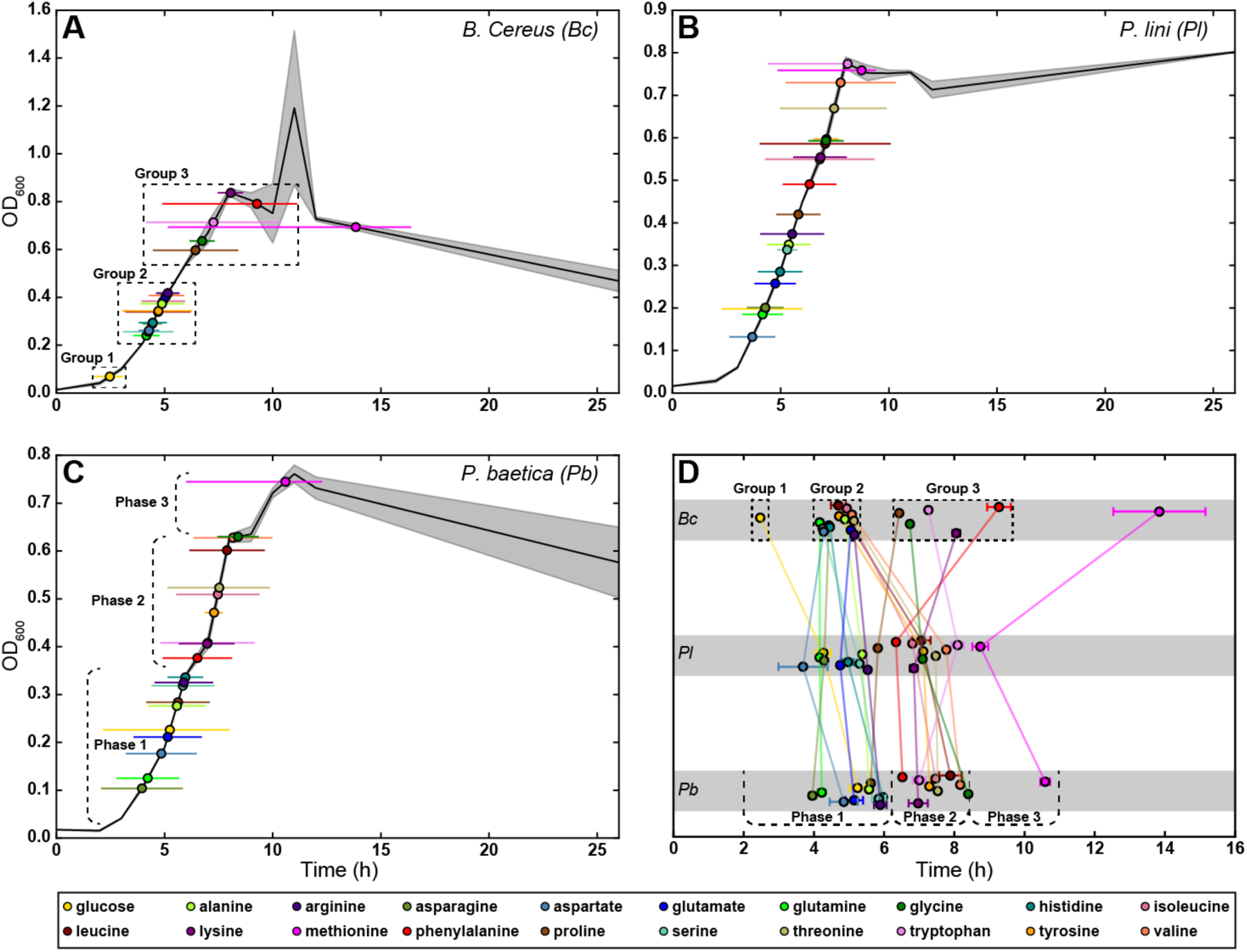
T_h_ and width for each compound mapped onto the growth curve of each strain. Colored circles represent average T_h_ and colored horizontal lines represent the average usage window (time of depletion from 90% to 10% of total resource used by the strain). Solid black line is the average OD600 of each strain measured over time (n=3), with shading representing standard deviation. **D)** Comparison of T_h_ values between strains, of all compounds, with error bars representing standard error. Dashed boxes in **(A)** and **(D)** indicate the grouping of compounds utilized by *Bc*, and dashed brackets in **(C)** and **(D)** indicate the different growth phases observed for *Pb*.

It is surprising that for these three species we observed three different combinations of growth curve and substrate utilization profile: a temporally distinct grouping of compound utilization with only one observed growth phase (**Figure 2A**), an even distribution of substrate utilization with only one growth phase (**Figure 2B**) and an even distribution over multiple growth phases (**Figure 2C**). This is quite significant given that two of the species belong to the same genus (*Pl* and *Pb*). This suggests that the metabolic regulatory systems between the two species are different: while *Pb* slows down its growth, presumably because it is undergoing a large-scale “switch” of metabolic systems, *Pl* does not, which may indicate that either all its metabolic systems are constitutively active, or the regulation of the systems is so perfectly timed that the organism can seamlessly switch from one metabolic regime to another. *Bc* may also have an efficient metabolic regulatory system, as even though we observe distinct temporal gaps between groups of compounds, we did not observe multiple growth phases.

To compare the differences in substrate depletion between species, we compared T_h_ across the three species (**Figure 2D** and **Supplemental Table 1**). Across all three species, glutamine, glutamate, alanine, arginine, proline, and asparagine, were half-depleted within one hour of each other. Additionally, the T_h_ values across all substrates for the two *Pseudomonas* species were close, but not identical, consistent with their short phylogenetic distance but different species identity (**Figure 2D**); a similar observation has been described previously ^22^. Considering the differences in growth curves between the two species, this is quite intriguing, as the general order in which the species consume the metabolites is not different, but there is this difference in growth profiles, supporting the hypothesis that there could be significant physiological differences between such closely related species.

*Bc* was markedly different from the two pseudomonads, differing greatly in the amount of time it depleted 8 of the compounds (**Figure 2D** and **Supplemental Table 1**). Of these, the utilization of glucose was particularly interesting, as it was predominantly depleted before there was any appreciable increase in biomass (**Figure 2A**). This may indicate that there is a significant delay in substrate conversion to biomass in this species, or that *Bc* rapidly transforms glucose into some other compound, for example glycogen.

We next wondered if the preferred substrates offer some physiological benefit over less preferable substrates. It is a general assumption in microbiology that substrates consumed first may be more advantageous than those consumed later ^24^, and that this would depend on the competitive ‘strategy’ of the organism. Major strategies suggested include maximal biomass production rate, maximal growth rate and maximal biomass yield. Generally, copiotrophs are thought of as r-strategists (maximal growth rate) and oligotrophs as K-strategists (maximum yield) ^25^,^26^. Given the relatively fast growth rates and high substrate concentrations in this study we would expect that the order of substrate consumption would be related to maximal growth rate or biomass production rate 27.

We tested some of these general assumptions by comparing the calculated T_h_ values and maximum usage rate of each compound to the specific growth rate, starting molarity of the compound, and predicted total protein composition of each species, in order to determine what the substrate preference order might be correlated with (**Figure 3** and **Supplemental Figure 3**). The specific growth rate of a species on a compound was determined by growing the species on that compound as a sole carbon source (see **Materials and Methods**). Surprisingly, the only significant (p < 0.05) correlations between all of these tests were that the specific growth rate of *Pl* on a given compound was weakly correlated with the T_h_ of that compound (r = −0.652, p = 0.030), and moderately correlated with the maximum depletion rate of that compound by *Pl* (r = 0.791, p = 0.004) (**Figures 3C**,**D**). These correlations support the common assumptions listed above, especially for flux balance analysis, as the compound that provides the higher rate of growth is depleted earlier and more rapidly than others. It is interesting that glucose did not confer the fastest specific growth rate for any of the strains, despite glucose generally being considered a superior source of energy. This is not surprising, however, as it is known that pseudomonads preferentially use amino acids over glucose ^28^. The rationalization of this phenotype is that in the soil environments where many pseudomonads (and *B. cereus*) live, decomposition products such as amino acids and organic acids are more readily available than sugars ^28^. However, the lack of any strong or significant correlations in the bacillus and the other pseudomonad indicates that there are other factors at play that determine an organism’s preferred substrate usage. It is apparent that not all microbes prefer to use substrates sequentially at all; the grouping of substrate utilization by *Bc* is a striking example of this. The resources within the second utilization group (**Figure 2A**) conferred a wide range of specific growth rates, from zero to the highest observed for all substrates, and all were utilized within two hours of each other (**Figure 3A**). It is likely the case that the simultaneous usage of these substrates confers the greatest physiological advantage. *Bc* could possess a metabolic strategy that does not perfectly follow the well-established paradigm of catabolite repression. Ultimately, it is clear that bacteria dramatically differ in regulation of catabolite uptake, and it is not prudent to make general assumptions on microbial metabolism based solely on observations from a few model organisms and/or the energetic potential of substrates.

**Figure 3.**
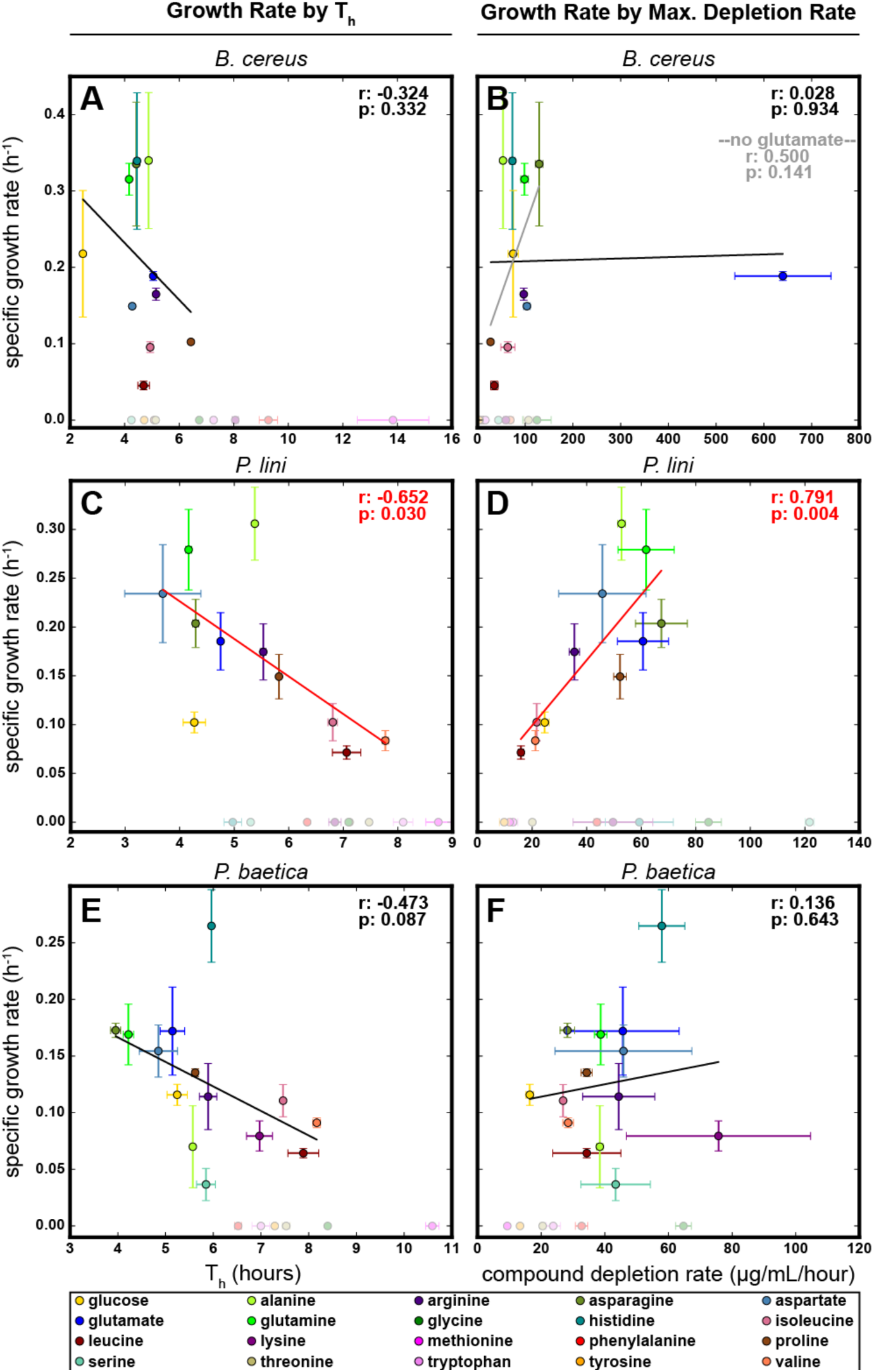
Correlations between specific growth rate on a compound as a sole carbon source, and T_h_ (**A, C, E**) or maximum compound depletion rate (**B, D, F**) in complete defined medium for species *Bc* (**A, B**), *Pl* (**C, D**), and *Pb* (**E, F**). Compounds that did not support growth as a sole carbon source (specific growth rate of zero) are shaded lighter at the bottom of each plot. Pearson correlation coefficients (r) and p-values (p) for the set of compounds for which the specific growth rate was nonzero are depicted in the upper-right of each plot. Correlations that had a p-value less than 0.05 were colored red. Error bars depict standard error.

Our experiments to test these correlations yielded a number of interesting results in addition to those described above. First, all three species grew on glucose as the sole carbon source without added amino acids. This was not predicted based on genomic predictions of the species in the Integrated Microbial Genomes (IMG) database (img.jgi.doe.gov), which indicated auxotrophy for lysine, phenylalanine, tyrosine, histidine and serine in the case of *Bc*, and for lysine, histidine, leucine, and coenzyme A for *Pl* and *Pb*. This observation highlights that all computational predictions should be treated as only suggestions, and should always be tested experimentally before making any assertions. Additionally, there were a number of compounds that did not support growth as sole carbon sources, but were depleted throughout the growth of the species in our complete defined medium (**Figure 3**, lightly shaded compounds). This finding indicates that caution should be employed when making physiological assertions based on single-substrate studies, for example those that have individual substrates arrayed in multi-well plates; many microbes can only utilize certain compounds when other substrates are present, the phenomenon of co-metabolism ^29^. We should note, however, that we do not know the details of how these compounds are depleted in the rich defined medium, only that they are depleted from the medium; they may simply be exogenously transformed. Finally, we observed the maximum depletion rate of all the substrates by the three species to be less than 130 μg/mL/hour except for glutamate depletion by *Bc*, which we calculated to be about 640 μg/mL/hour (**Supplemental Table 1**). This rate corresponds to a near instantaneous depletion of glutamate by *Bc* at about 5 hours into the growth curve (see **Supplemental File 1**), which is towards the end of the second group of compounds utilized by this species (**Figure 2A**). Why glutamate would be depleted so much faster than the other compounds for *Bc* is a mystery, but it does suggest that there is something unique about the compound that requires or allows for the flux to be so rapid. Interestingly, in a previous study of metabolite depletion of a mixture of 470 compounds glutamate was one of two metabolites depleted by all of the isolates ^14^, so it is clearly an important or high-value compound that *Bc* may have evolved to deplete quickly in order to gain a competitive advantage.

### Predicting consortium metabolism based on models of individual isolates

Having modeled the substrate usage of each species for each compound, we hypothesized that these models could be combined to predict how a consortium composed of the three species might utilize the substrates. We simulated the time-dependent depletion of each compound by a consortium composed of the bacillus and two pseudomonads (see **Materials and Methods**, **Equation 2**, and **Algorithm 2**). Briefly, the functions describing the compound usage by each species were summed (**Supplemental Figure 2A**), and the time at which this summed use curve reached the total available compound was determined. This time of depletion was then used to predict how much of a given metabolite each species would have utilized when grown in co-culture, and the compound usage by each species was re-modeled (**Supplemental Figure 2B** colored dashed lines) and added together to form the co-culture prediction (**Supplemental Figure 2B** solid black line). These predictive models allowed us to make several hypotheses that are relatively simple to test. First is the usage curve of each metabolite by the co-culture. Related to this, we can predict the time at which all of a given metabolite will be depleted, and when all metabolites will be depleted. From this we predict that 14 compounds will be nearly depleted (less than 10% of starting concentration) by six hours, and all but methionine will be completely consumed by 9 hours (**Figure 4**). Based on this, one could reasonably argue that a consortium composed of these three species would reach stationary phase sometime between 6 and 9 hours, in contrast to the individual species, which all reached stationary phase after 9 hours.

**Figure 4.**
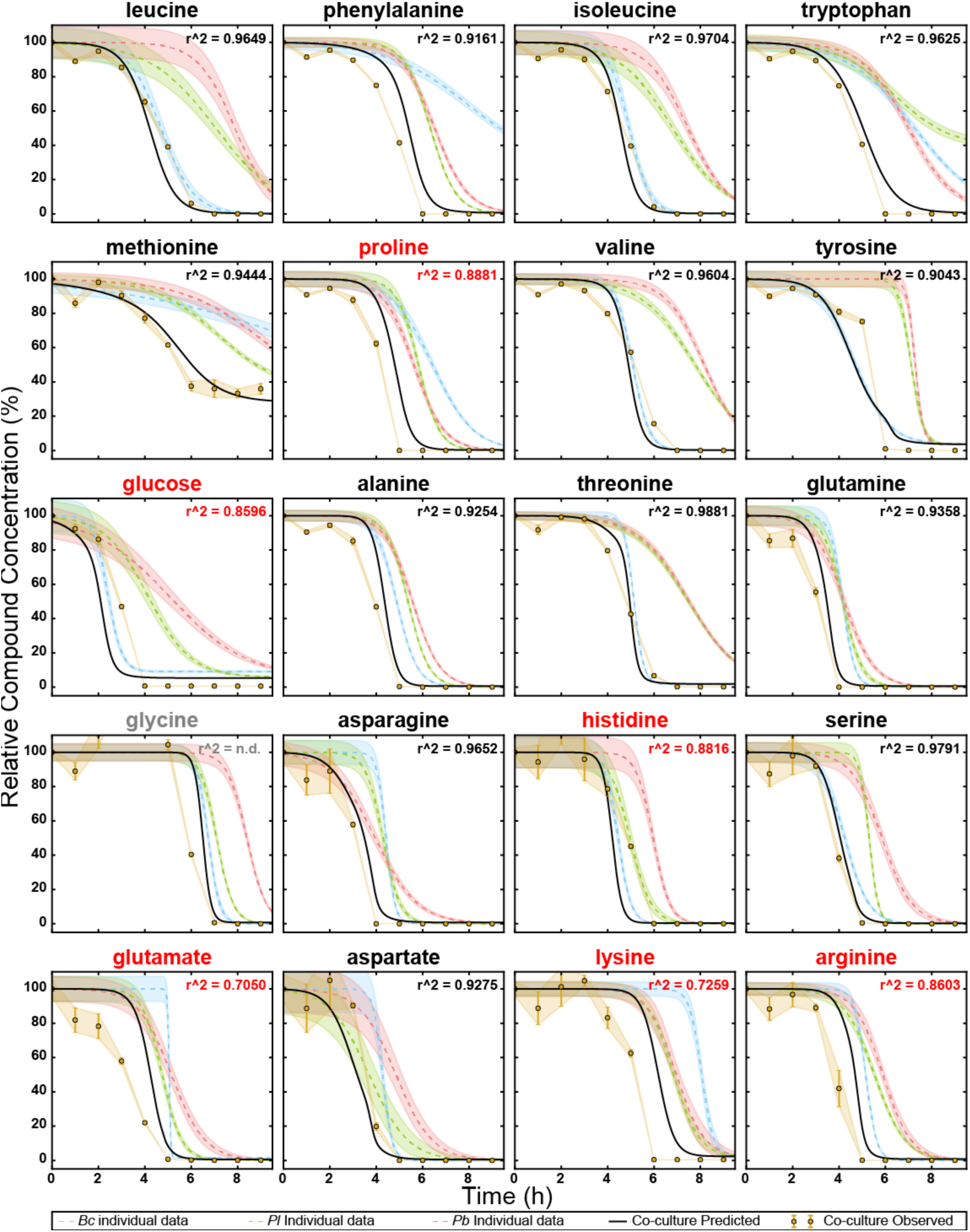
Co-culture observations compared to predictions, normalized to t0 concentration of each metabolite. Blue, green, and red dashed lines represent the observed depletion of each compound by *Bc, Pl*, and *Pb*, respectively, when grown in isolation. The solid black line is the predicted depletion of a co-culture of all three strains. The golden circles represent the measured compound concentration in the co-culture medium. Error bars and/or shading represent standard error (n=3). Glycine at time point 4 could not be calculated because the measurement was outside the dynamic range of the calibration curve, and the r^2^ was not determined (n.d.) for glycine. Non-normalized figure is shown as **Supplemental Figure 5**.

To test our predictions, we inoculated a 3-member co-culture at equal optical density in the defined medium (see **Materials and Methods**), collected supernatant time points every hour, and measured the concentrations of all 20 substrates as described for monocultures. We found that many of our predictions were valid: nearly all compounds (17) were depleted to below 10% of starting concentration by 6 hours (**Figure 4**, gold), and the co-culture accordingly reached stationary phase at this time as well (**Supplemental Figure 4**), presumably because all available substrates were consumed.

### Compounds that follow the model are evenly shared

When analyzing the kinetics of depletion of the compounds, we observed that many (13) compounds agreed very well with the prediction, having R^2^ values greater than 0.9 (**Figure 4**). Most of the compounds with high R^2^ values began to decrease slightly earlier or at a slightly faster rate than predicted, which could be attributed to experimental error in initial culture density. However, the depletion of most compounds were still very close to the predicted model, indicating that the shared usage between the species could be very close to “blind” conditions, where the presence of other species does not affect the substrate usage decisions of each individual species. It is important to note that the high substrate concentrations likely explain the successful predictions using this simple modeling approach. Specifically, the substrate concentrations, initially at high micromolar concentrations, are likely well above the K_m_ for the transporters and rate-limiting enzymes. For example many bacterial amino acid transporters have K_m_ values in the low micromolar range ^30^,^31^, such that the transporters and enzymes are saturated. We anticipate that much more detailed models accounting for substrate concentration would be required at soil-and groundwater-relevant substrate concentrations, which can be as low as 0.5-10% of the concentrations used in this study (^32^ and Jenkins et al., in preparation).

### Compounds that deviate from the model

The remaining 7 compounds (glucose, histidine, glutamate, lysine, arginine, proline, and glycine) deviated significantly from our predictions (R^2^ < 0.9) (**Figure 4**, red text), suggesting some additional species-species interaction(s) is/are present that affect the depletion of those compounds. It is intriguing that we detected metabolites that showed both positive and negative deviations.

Glucose and histidine were both depleted more slowly than predicted. The simplest explanation for this is that the metabolic systems that deplete these compounds are indeed concentration dependent. Another possibility for this would be that there is a buildup of product in the co-culture that exerts feedback inhibition on the metabolism of these two compounds. This is easily rationalized for histidine utilization, which is an expensive process for bacteria ^33^; they may be exposed to better carbon sources in a mixed culture as a byproduct of another microbe. However, glucose being utilized slower is curious. In the monoculture experiments, we observed *Bc* to deplete glucose before it or either pseudomonad even started producing appreciable biomass (**Figure 2**). Perhaps this behavior is inhibited in the presence of the pseudomonads or is a result of changes in the community structure over the experiment, the assessment of which are unfortunately beyond the scope of the current study.

In contrast, glycine, proline, lysine, arginine, and glutamate were all depleted faster than predicted. This is more difficult to explain and suggests at least one microbe has altered its phenotype due to the presence of other microbes, or that other exometabolites are influencing consortial behavior. For example, one species may have up-regulated metabolic pathways involving these compounds in an effort to outcompete others, either for the purpose of direct competition for the substrate, or in order to synthesize antibiotic compounds ^34^. Alternatively, another member may have otherwise sequestered those compounds, effectively taking them out of a common pool, for example by converting the compound into some storage molecule, or sequestering it in a way similar to how siderophores sequester iron. Testing these hypotheses would require an extensive untargeted metabolomics study, an extremely interesting direction for future studies. Another potential reason for this early depletion is that the co-culturing of these microbes has resulted in an emergent function of increased flux of the substrate(s) through the system. This could be due to a cross-feeding effect where one microbe depletes an inhibitory compound of another microbe or one microbe’s products induce the co-metabolism of that product and one of these substrates.

## Conclusions

This study examining substrate competition for 20 abundant substrates by 3 species demonstrates that at least some portion of the metabolic behavior of a microbial consortium can be predicted by measuring the metabolism of microbes grown in monoculture. This likely can also apply to more complex situations, for example separately measuring the metabolism of an existing microbial community and a foreign isolate, and predicting what the metabolic function might be if the isolate were introduced into the community. In any system, compounds that do not fit the predictions indicate emergent functions of the coculture and may highlight substrates that are somehow affected by species-species interactions. These may be occurring passively in the cases of feedback inhibition and co-metabolism, or actively in the case of one species altering its phenotype in order to outcompete others. Further studying these outlier substrates can shed light on metabolic interactions between microbes within a community. Ultimately, incorporating this predictive strategy when studying community metabolisms can help pinpoint interesting biological questions, as well as aid in the design of synthetic consortia.

## Materials and Methods

### Isolates and identification

The 16S rRNA gene for each isolate was amplified using primers 27F (AGAGTTTGATCMTGGCTCAG) and 1492R (CGGTTACCTTGTTACGACTT), and sequenced at the Eurofins sequencing facility (Eurofins MWG Operon LLC, Louisville, KY). Forward and reverse sequences were manually merged and used as queries using nucleotide BLAST against the 16S rRNA sequence database at NCBI.

### Phylogenetic Tree Construction

16S rRNA gene sequences were obtained from IMG (img.jgi.doe.gov), except for *B. cereus*, *P. lini*, and *P. baetica*, which were directly sequenced (see above). Gene sequences were aligned using MUSCLE ^35^,^36^, curated using GBlocks ^37^, and the tree was constructed using PhyML ^38^ with 100 bootstraps, using the phylogeny.fr web server ^39^,^40^. The final tree was rendered using FigTree (http://tree.bio.ed.ac.uk/software/figtree/).

### Growth medium and culturing

All bacterial species were initially inoculated from frozen glycerol stocks onto an R2A agar plate prepared using Difco R2A Agar (BD, Franklin Lakes, NJ) and incubated overnight at 30 °C. The medium used for metabolomics experiments consisted of 1x Wolfe’s vitamins and 1x Wolfe’s minerals solutions ^41^, 1.5 mg/mL ammonium chloride, 0.6 mg/mL potassium phosphate, and 0.1 mg/mL each of D-glucose and the following L-amino acids: alanine, aspartate, glutamate, phenylalanine, glycine, histidine, isoleucine, lysine, leucine, methionine, asparagine, proline, glutamine, arginine, serine, valine, threonine, and tryptophan. Tyrosine was additionally supplied at 0.01 mg/mL. Species were individually cultured in 5 mL of this medium overnight at 30 °C from the R2A plate, then washed 3x by centrifugation at 5,000 xg and resuspending in fresh medium. Washed cells were used to inoculate 50 mL of the medium in 250 mL Erlenmeyer flasks, at an initial optical density (OD_600_) of 0.012-0.017 as measured by a SpectraMax Plus 384 plate reader. These cultures were incubated at 30 °C, shaking at 200 rpm. For co-culture experiments, 50 mL cultures were inoculated with an OD_600_ of 0.012 of each species, resulting in an initial co-culture density of 0.036. 200 μL of cell culture was aspirated for OD_600_ measurements taken in a 96-well Falcon tissue culture plate with flat bottom. For all growth experiments, the water used to prepare the medium and uninoculated medium were incubated alongside the experimental flasks, as controls.

Growth assays of species on individual carbon sources were performed in 96-well Falcon tissue culture plates with flat bottom and low evaporation lid, in a total volume of 200 μL. The medium consisted of the same concentrations of Wolfe’s vitamins and minerals, ammonium chloride and potassium phosphate. Individual carbon sources were added at a concentration of 0.5 mg/mL. Species were pre-cultured and washed as before, and wells were inoculated at an OD_600_ of 0.05. The plates were incubated at 30 °C, shaking at “medium” speed in BioTek Synergy HT and Tecan Infinite F200 Pro plate readers, for 48 h.

### Metabolomics sample extraction

Hourly time points of 1 mL of cell culture and controls (see above) were aspirated and centrifuged at 5,000 xg to pellet the cells. 800 μL was aspirated from the top, taking care not to disturb the cell pellet, and split into two 400 μL aliquots, which were immediately frozen at −80 °C. A calibration curve was created with the medium used for culturing: 1x culture medium, 1/2x, 1/10x, 1/100x, 1/1000x, and 1/10000x dilutions were prepared using culture medium without any carbon sources as the diluent. All experimental, control, and calibration curve samples were lyophilized overnight, and metabolites were extracted in 300 μL methanol with 25μM ^13^C-phenylalanine for use as an internal standard. Final extracted samples were stored in Agilent 96-well sample plates and immediately analyzed via LCMS or stored at −80 °C.

### Metabolomics data acquisition and quantification

An Agilent 1290 LC system equipped with a ZIC-pHILIC column (150 mm × 2.1 mm, 5 μm 100 Å, Merck SeQuant) was used for metabolite separation with the following LC conditions: solvent A, 5 mM ammonium acetate; solvent B, 9:1 acetonitrile:H_2_O with 5 mM ammonium acetate; flowrate: 0.25 mL/min; timetable: 0 min at 100% B, 1.5 min at 100% B, 25 min at 50% B, 26 min at 35% B, 32 min at 35% B, 33 min at 100% B, and 40 min at 100% B; column compartment temperature of 40 °C. Mass spectrometry analyses were performed using Agilent 6460 triple quadrupole mass spectrometer. Agilent software (Santa Clara, CA): Optimizer was used for establishing fragmentor and collision cell voltages as well as precursor and product ion transitions while Mass Hunter QQQ Quantitative Analysis (version 6.0) was used for compound quantification. Retention times, collision energies, and transitions for each compound are listed in **Supplemental Table 2**.

### Substrate depletion modeling

The Anaconda package and IPython notebooks were used for all computational tasks ^42^, which will be made publicly available at https://github.com/biorack in the “Predicting metabolic properties of a microbial co-culture” repository upon manuscript publication by a peer-reviewed journal. Data were stored and organized using Pandas ^43^ and NumPy ^44^, and graphs created using Matplotlib ^45^. Metabolite depletion was modeled using **leastsq** from **scipy.optimize** ^46^, fitting the data to the Behrends model (eq 1):

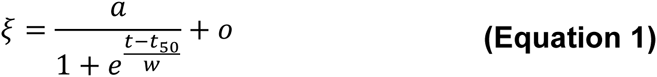

Where *a* is amplitude and *o* is offset (see **Figure 1**). These two parameters were defined from the data: amplitude was defined to be the average of the t=0 data point and the maximum value data point in the data set of each compound, and offset was defined as the lowest value in the data set. All other parameters were solved using **leastsq**, with the criteria that they had to be positive values. The exact steps are shown in **Algorithm 1**:

#### Algorithm 1: modeling depletion of each substrate by each species

1 *species* ← the set of species used in the experiment

2 *substrates* ← the set of compounds measured in the experiment

3 *t* ← time dimension of the experiment

4 **for** *i* **in** *species*:

5 **for** *j* **in** *compounds*:

6 *data*_*ij*_ ← measurement series of *substrate j*

7 *o*_*ij*_ ← **minimum**(*data*)

8 *a*_*ij*_ ← **average** (*data*_*ij*_[0] and **maximum**(*data*_*ij*_))

9 **leastsq** parameter fitting of *t*_50_*ij*__ and *w*_*ij*_ to *data*

10 
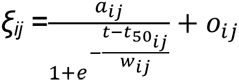

T_h_ and usage window values were calculated from the Behrends model. All correlation coefficients and p-values were calculated using the **pearsonr** function in the **stats** package of **scipy**.

### Co-culture predictions

The equations representing the depletion of a compound by a species were subtracted from the initial starting concentration of the compound, creating an expression that represented the amount of compound used by each species over time; these are the curves shown in **Supplemental Figure 2A**. These expressions were summed to generate an approximate total usage curve, and the time at which this curve crossed the total amount of available compound was determined. The amount of available compound was defined to be the starting concentration of a compound minus the lowest offset parameter between the three species, as the species with the lowest offset parameter for a substrate will presumably deplete the substrate to that level, but not more, even in a co-culture. The time of total depletion was used to approximate the amount of compound that each species would have consumed by that time. The individual usage curves were capped at this compound level at this time, and transformed back to compound depletion curves, which were then used to re-fit to the Behrends equation, generating new models of compound depletion in mixed conditions. These new models were then summed, producing the predicted total co-culture usage of each compound. This can be summarized by the general **Equation 2**:

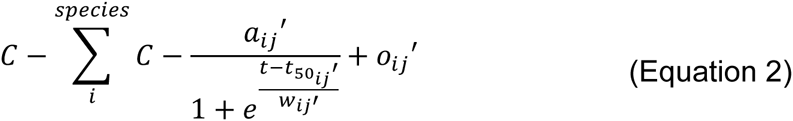

Where C is the total amount of substrate *j* that is available to the mixed culture of set *species*. This is defined as the starting concentration of *j* minus the smallest *o*_*j*_ in *species. a*_*ij*_′, *o*′_*ij*_, *t*_50_*ij*__, and *w*_*ij*_′, are parameters that describe the depletion of *j* by species *i* in the co-culture of the individual in the set *species*, shown in **Algorithm 2**:

#### Algorithm 2: Predicting co-culture substrate usage

1 **for** *j* **in** *substrates*:

2 *o*_*j*_’ ← minimum (*o*_*j*_)

3 *C* ← starting concentration of substrate *j* **minus** *o*_*j*_’

4 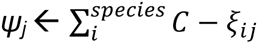

5 *t*_*d*_*j*__ ← *t* when *ψ*_*j*_ = *C*

6 **for** *i* **in** *species*:

7 *Φ*_*ij*_ ← *ξ*_*ij*_(*t*_*d*_*j*__)

8 *temp* ← *ξ*_*ij*_(*t*_*d*_*j*__:*t*_*n*_) = *Φ*_*ij*_

9 *o*_*ij*_’ ← *ξ*_*ij*_(*t*_*d*_*j*__)

10 *a*_*ij*_’ ← starting concentration of substrate *j* minus *o*_*ij*_’

11 **leastsq** parameter fitting of *t*_50_*ij*__′ and *w*_*ij*_’ to *temp*

## Acknowledgements

The strains used in this study were a generous gift from Romy Chakraborty at Lawrence Berkeley National Laboratory.

This material by ENIGMA-Ecosystems and Networks Integrated with Genes and Molecular Assemblies (http://enigma.lbl.gov), a Scientific Focus Area Program at Lawrence Berkeley National Laboratory is based upon work supported by the U.S. Department of Energy, Office of Science, Office of Biological & Environmental Research under contract number DE-AC02-05CH11231

